# A sex-balanced longitudinal developmental task-free fMRI rat dataset

**DOI:** 10.64898/2026.06.04.730202

**Authors:** Daniel McLoone, Andrew Breen, Claire McParland, Andrew Harkin, Clare Kelly

## Abstract

Here we present a preclinical longitudinal developmental dataset spanning pre-puberty (juvenile) to early adulthood in Wistar rats. Thirty-six rats (18 female) were scanned during five sessions at approximately postnatal day 28, 35, 49, 70 and 91. A standardised consensus protocol was used for animal sedation and MRI data acquisition (structural MRI and resting-state fMRI). The dataset also includes daily body weight measurements, day of puberty onset, and daily cytological smear images for oestrous tracking. This openly available dataset addresses the need for sex-balanced developmental preclinical datasets spanning puberty and enables researchers to investigate longitudinal brain functional developmental trajectories and Sex As a Biological Variable (SABV).

## Background & Summary

Adolescence is a period of major transition from childhood to adulthood. It is characterised by physical and sexual maturation^1^, profound cognitive and emotional changes, and an increasingly complex social environment^2–4^. During puberty, sex hormones begin developmentally-timed, recurrent interactions with the brain, appearing to ‘reorganise’ brain and behaviour^5^, with sexual dimorphisms in many of these changes^6–8^. In humans, these changes are “entangled” with psychosocial dynamics related to gender and its development^9^.

Adolescence represents a sensitive period of neuroplasticity, during which cognitive, emotional, and social capacities^8,10–13^ mature and brain structure and function are significantly altered^3,14,15^. This critical period of plasticity provides an important developmental window of opportunity for learning and thriving, but also constitutes a period of heightened risk, with many mental disorders emerging after the onset of puberty^2,16^. The relationship between adolescence and mental illness is complex and multidirectional, involving biology, environment, and psychosocial factors, but it appears that the plastic pubescent brain is especially vulnerable to stress-induced developmental perturbations, which can manifest later as psychiatric disorder^17–19^. The postpubertal emergence of salient sexual dimorphisms in the prevalence of some psychiatric disorders^20–22^ further underscores a likely role for pubertal processes, such as the actions of gonadal hormones, in adolescent vulnerability to mental health difficulties.

Given the significance of the adolescent transition for long-term outcomes, considerable research effort has been dedicated to charting trajectories of adolescent brain development using magnetic resonance imaging (MRI) methods. One goal of this research is the delineation of growth curves for brain development, analogous to those for height, weight, and head circumference, which have served as medicine’s gold standard for spotting problems in development for over a century. Evidence of deviation from growth curves for neurocognitive development could support the early identification of at-risk individuals, allowing us to intervene to divert developmental trajectories away from illness towards health^23,24^. Collaborative retrospective data pooling efforts have demonstrated the feasibility and potential value of charting growth trajectories for brain structure^25^ and function - using resting-state functional MRI (rs-fMRI) analyses that map the brain’s intrinsic functional architecture^26^. Large-scale prospective human studies, such as the ongoing Adolescent Brain Cognitive Development study (ABCD)^27^ and Lifespan Chinese Color Nest Project^28,29^ are directly aimed at providing the kinds of rich longitudinal neuroimaging, cognitive-behavioural, and clinical data that will enable the characterisation of growth curves for brain function. Critically, longitudinal designs allow for more powerful developmental analyses capable of dissociating aberrant or delayed maturation from individual differences such as sex, better supporting causal inferences about sources of influence on brain development^30,31^.

However, an inherent challenge for human longitudinal studies is the long timescale over which human development unfolds. Human adolescence spans more than a decade - as a result, human longitudinal studies require careful planning, considerable resources, and significant commitment from both researchers and participants^32^. Human longitudinal studies are therefore costly and unavoidably prone to attrition, while also exposed to a range of uncontrollable environmental influences that hamper causal inferences.

Although preclinical research cannot capture the full complexity of human brain, cognitive, and socioemotional development, longitudinal animal models nonetheless offer the advantage of avoiding these specific pitfalls. Rodents have a far shorter adolescence than humans, reaching adulthood in less than three months. Rodent models therefore reduce resource and time burdens while enabling collection of longitudinal data points under conditions of strict environmental control, minimising sources of extraneous influence.

Rodent models also permit direct experimental manipulations (e.g., controlled exposure to stress) that are not possible in humans, enabling causal inferences. These factors motivated the current open dataset, which comprises rs-fMRI data collected from 36 Wistar rats, scanned longitudinally at five timepoints spanning the juvenile (pre-puberty) period to early adulthood. Importantly, rs-fMRI studies have demonstrated that functional brain networks are conserved across species^33–35^; application of the same rs-fMRI developmental mapping methods routinely applied in humans therefore offers a translational bridge from the mechanistic and causal insights obtainable only in animal models. To maximise the potential future use of our data and generalisability of findings, we used a recently developed standardised consensus protocol for animal sedation and MRI data acquisition (structural MRI and resting-state fMRI)^36^. Data are shared in raw as well as fully preprocessed form, with timeseries derivatives, to maximise ease of use and minimise unnecessary duplication of effort and use of computational resources.

One shortcoming of rodent neuroscience studies is that they have historically tended to exclude female rodents, due to assumptions of increased hormonal and behavioural variability^37,38^. Recent research suggests that male rodents may display just as much variability as females, however^39^. Sex-blindness has likely impeded progress by reducing the translational validity of preclinical research, since findings obtained in one sex may not translate well to the other, particularly when they must also be translated across species. In recognition of the impact of sex-blindness on scientific knowledge, reproducibility, and equity, funding bodies such as the NIH and Horizon Europe now require the incorporation of Sex as a Biological Variable (SABV) in research, although many challenges and questions remain^40,41^. In providing a sex-balanced rodent dataset, the current dataset upholds the principle of *Sex As A Biological Variable* ^42^. Data on timing of puberty onset and daily vaginal smear images (for oestrous tracking) are shared, to allow for more sophisticated analyses that take into account pubertal/oestrous status.

In sharing these data, we provide a dataset that can be used by future researchers to answer a wide range of translational questions related to brain development during the adolescent transition. Our dataset also addresses the urgent need for sex-balanced preclinical data, and upholds 3Rs principles^43^, by enabling further research to be conducted without the need for additional animals.

## Methods

### Animals and study design

Male and female Wistar Han rats were obtained from Inotiv (United Kingdom) and were bred within the Comparative Medicine Unit in Trinity College Dublin. Rats were weaned on postnatal day 21 (P21), with the day of birth considered P0. Animals were housed in pairs in individually ventilated cages (Innovive Innocage® Tall IVC Rat), on a 12-hour light-dark cycle with ad libitum access to food and water. Each cage had nesting material, a red polycarbonate tunnel and a red tall rat loft. Animals were kept in climate-controlled rooms set to 22°C (±2°C) with relative humidity levels of 50%. Two male experimenters handled and carried out all experimental procedures on the rats, while staff providing general animal care (water, bedding, etc.) were both male and female.

Animals were handled daily, with puberty onset recorded, and the oestrous cycle tracked in females once puberty was reached. In females, puberty onset was defined as vaginal opening^44^, while in males, balanopreputial separation was used^45^. To track oestrous a sterile cotton swab was moistened with sterile saline and any excess flicked off. The swab was inserted approximately 1cm into the vaginal canal and rotated one revolution. Swabs were then rolled onto a glass slide and allowed to dry. Slides were stained using Wright Stain, Modified (WS16, Sigma-Aldrich Gillingham). Using a dropper, slides were covered in stain for one minute. Next an equal volume of deionised water was added and mixed by gently swirling for three minutes. Slides were then washed with deionised water and allowed to dry. Slides were imaged at 20x using a light microscope (Olympus BX51) equipped with a digital camera (Olympus DP23).

Figure 1 shows the scanning sequence for each data collection wave. Animals underwent five MRI scans at approximately P28, P35, P49, P70 and P91 (see participant.tsv and subject specific session.tsv for age-at-scan information along with each animal’s date of birth, sex, litter identifier, experimental wave, and date of puberty onset). Scans were acquired over four waves, with 4 to 12 animals per wave.

**Figure 1.**
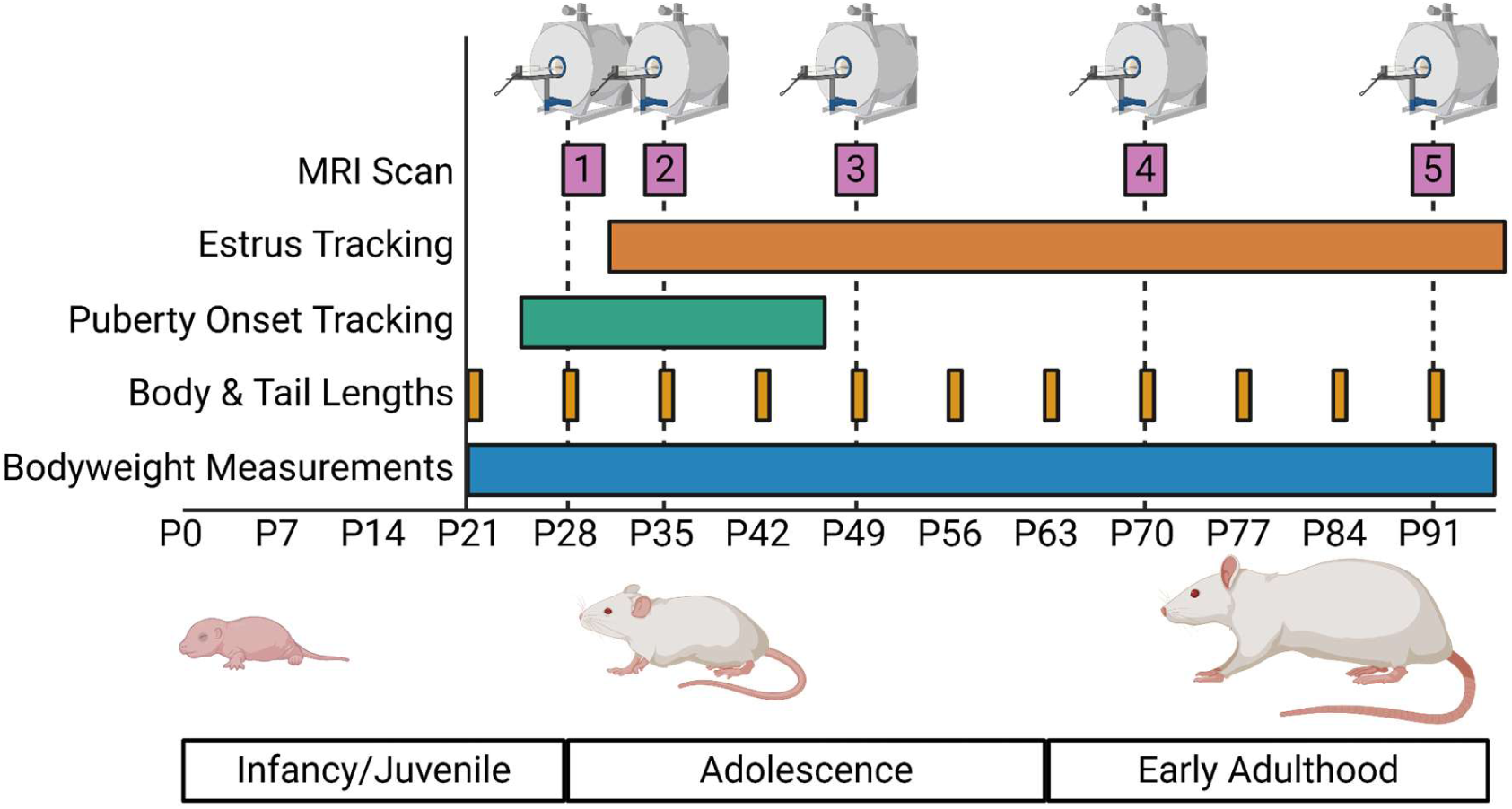
Experimental timeline and data collection. The day of birth was considered P0. Animals were weaned on P21 and weighed daily starting on P21. Body and tail lengths were measured weekly starting on P21. Puberty onset tracking began at P26 and continued until puberty onset. Oestrous tracking began in females following puberty onset, and continued daily. Resting-state MRI scans were acquired at approximately P28, P35, P49, P70 and P91.Created in BioRender. Mc Loone, D. (2026) https://BioRender.com/wtr93yk

At the end of data collection (approximately P91) rats were deeply anaesthetised with sodium pentobarbital (100 mg/kg i.p.), with depth of anaesthesia determined using the toe pinch method. Rats were then transcardially perfused with cold phosphate buffered saline (PBS), followed by 4% paraformaldehyde (PFA). Following perfusion, animals were decapitated and their brains removed. Brains were then placed in 4% PFA and stored overnight at 4°C. Next, brains were cryoprotected in 30% sucrose solution and stored at 4°C for 72 hours or until brains sank to the bottom of the tubes. Brains were then placed in isopentane on dry ice and snap frozen and stored at -80°C to allow for future post-mortem research.

### Ethics statement

All animal experiments and procedures were conducted in compliance with the European Directive 2010/63/EU on the protection of animals used for scientific purposes, approved by the Animal Research Ethics Committee in Trinity College Dublin and carried out under licence granted by the Health Products Regulatory Authority (project authorisation AE19136/P173).

### Neuroimaging data collection

A Bruker 7T BioSpec 70/30 USR system (Bruker, Ettlingen Germany) equipped with a 1H receive-only 8 x 1 rat surface array coil, 72mm transmit coil, BGA9SHP gradients and Paravision 7 software was used for all imaging sessions. The StandardRat protocol was used for both anaesthesia and scan parameters (https://github.com/grandjeanlab/StandardRat);^36,46^. Anaesthesia induction was carried out using 4% isoflurane in 1000ml/min oxygen and 400ml/min nitrogen. This was followed by a bolus of medetomidine (0.05mg/kg, s.c.). Animals were maintained at 0.4% isoflurane in 200ml/min oxygen and 400ml/min nitrogen, with a continuous infusion of medetomidine (0.1 mg/kg/h, s.c.) starting 15 minutes after the bolus (NE-300 syringe pump, New Era Pump Systems, Inc, New York, USA). The EPI scan began 40 minutes after the bolus of medetomidine had been delivered. Isoflurane concentration was set to 0.4% at least 10 minutes before starting the EPI sequence. Once complete, the continuous infusion was stopped and atipamezole (0.5mg/kg, s.c.) was delivered to reverse the effects of medetomidine. Respiration and temperature were monitored throughout the scan using a respiratory pillow and rectal probe (Model 1025T, SAII, New York, USA), with body temperature maintained at 37°C using a water heating system connected to the MRI cradle.

For the anatomical scan, a T2w turboRARE sequence was acquired: repetition time (TR) = 2500ms, echo time (TE) = 30ms, RARE factor = 8, 18 interleaved axial slices of 1 mm with 0.1-mm gap, FOV = 25.6mm x 25.6mm, and matrix size of 128 x 128^36^.

For the functional scan, a gradient echo-echo planar sequence was acquired using parameters: TR = 1000ms, TE = 17ms, 18 interleaved axial slices of 1 mm with 0.1-mm gap, FOV = 25.6mm x 25.6mm, and matrix size of 64 x 64^36^.

### Neuroimaging data processing

Anatomical and functional scans were converted from raw Bruker format to Neuroimaging Informatics Technology Initiative (NIfTI) format and structured in Brain Imaging Data Structure (BIDS) format using BrkRaw v0.3.11 (https://github.com/BrkRaw/brkraw). All scans were reoriented to Right, Anterior, Superior (RAS) using Analysis of Functional NeuroImages (AFNI). All AFNI (afni/afni_make_build:AFNI_25.2.11) and Advanced Normalization Tools (ANTs) (antsx/ants:2.6.2) commands were run in docker containers (Docker version 28.3.3, build 980b856).

### Unbiased dataset template generation

Anatomical image masks were generated using Brain Extraction Net (BEN), a deep neural network for brain extraction^47^. BEN was run in an apptainer container available at https://github.com/grandjeanlab/apptainer/tree/e35a9dab1f972cbff03d3c59a41284c88602e0 3c/ubuntu_ben. One of the available weights for BEN (Rat-T2WI-9.4T) was used as the starting point for domain adaption as it was trained on T2w scans with similar parameters to those of StandardRat. To ensure the network was generalisable across different ages, sexes and experimental runs, all five developmental scans from one male and one female rat from each experimental wave were selected for training. Masks were manually created for each resulting in a training set consisting of 40 anatomical images and accompanying masks for domain adaption. The new updated weight generated was then used as the weight for inference to create masks for all 180 anatomical scans in the dataset. The masks created were then postprocessed by using ImageMath FillHoles and GetLargestComponent in ANTs.

DenoiseImage was applied to all anatomical images. Next a tissue mask was generated by ThresholdImage and N4BiasFieldCorrection was applied to voxels inside this mask in ANTs. To centre the brain in the FOV, anatomical images were cropped by 50 voxels from the inferior of each image using 3dZeropad in AFNI. Then, to improve the results from subject average template construction, the images were padded by adding 2 slices to both the anterior and posterior of each image using 3dZeropad in AFNI. The cropping and padding steps were repeated for the anatomical image masks.

To generate an unbiased dataset template, optimized_antsMultivariateTemplateConstruction, a re-implementation of the ANTs template construction pipeline was used https://github.com/CoBrALab/optimized_antsMultivariateTemplateConstruction^48^. The twolevel_modelbuild script was run in an environment with ANTs 2.6.2 and qbatch 2.3.1. For a given subject all inputs were aligned using their centre-of-mass before averaging. Four iterations of model building were used with SyN-control = 0.2,3,0.5, SyN-metric CC[2], convergence = 1e^-9^, with reuse affines and rigid update enabled. This resulted in 36 unbiased subject-average templates (first-level). These subject averages were then used to generate an unbiased population-average template (second-level). Padding added during template construction was then removed prior to preprocessing using ImageMath PadImage to reduce the image size by 18 voxels in all directions. The resulting unbiased population-average template was resampled to 0.2x0.2x0.2mm and used as the brain template for preprocessing.

### Functional data preprocessing

To centre the brain in the FOV, functional images were cropped by 25 voxels from the inferior of each image using 3dZeropad in AFNI. The preprocessing was conducted using the open-source Rodent Automated Bold Improvement of EPI Sequences (RABIES) software in a docker container (ghcr.io/cobralab/rabies:0.5.5; https://github.com/CoBrALab/RABIES)^49^. Subject anatomical scans were corrected for inhomogeneities. RABIES was run on each subject-wise to enable generation of an unbiased subject average template, which acted as an intermediate registration target for each session’s anatomical scan. To correct for EPI susceptibility distortions, the EPI was subjected to inhomogeneity correction, and then registered using a nonlinear registration to the anatomical subject-average for that subject. Finally, after calculating the transformations required to correct for head motion and susceptibility distortions, transforms were concatenated into a single resampling operation which was applied at each EPI frame, generating the preprocessed EPI timeseries in native space at a voxel resolution of 0.3x0.3x0.3mm.

After the first step in RABIES was carried out, the bold mask generated during preprocessing was replaced with a manually generated mask using 3dAutomask which better represented the brain coverage of each individual functional scan. Confound correction was then carried out in RABIES on the native space EPI timeseries. Voxelwise detrending was applied followed by bandpass filtering (0.01-0.1Hz). Thirty seconds was removed from each edge of the timeseries to account for potential edge artefacts following filtering. Next, 6 rigid motion parameters and the global signal were modelled at each voxel and regressed from the data. Finally, a spatial Gaussian smoothing filter was applied at 0.5mm full-width at half maximum. The confound corrected NIfTI images were then resampled to common space for technical validation.

Visual inspection was performed on preprocessing outputs for all scans for quality control. Eleven scans were excluded due to failing preprocessing: sub-2023080701_ses-2, sub-2023090104_ses-2, sub-2024011405_ses-1, sub-2024011406_ses-1, sub-2024011406_ses-2, sub-2024011501_ses-1, sub-2024011503_ses-1, sub-2024011504_ses-1, sub-2024011505_ses-1, sub-2024021401_ses-1.

For completeness, and to allow users to make their own choices regarding exclusion, the published dataset retains all (179) functional scans.

### Data Record

Data records are organised according to the Brain Imaging Data Structure (BIDS) standard (Version 1.8.0). The dataset is publicly available on OpenNeuro: ds007859; https://openneuro.org/datasets/ds007859/

The main folder contains a sourcedata folder, derivatives folder and 36 subject folders in BIDS format, with accompanying information including a dataset_description.json, participant.json, participant.tsv and README.md. The participants.tsv file contains each animal’s date of birth, sex, litter identifier, experimental wave and date of puberty onset. Each subject folder contains five sessions, with an anat and func folder for each. The subject level session.tsv file (sub-<ID>_session.tsv) provides the exact age and date for each scan and mean respiration for that subject. Exact dates have been provided to allow cross referencing with oestrous smear images.

The sourcedata folder contains an oestrous_smears folder and a growth_metrics folder. The estrus_smears folder has a subject folder for each female rat. Smears are labeled with the date of the swab and magnification of the image: sub-<ID>_date-<DATE>_acq-20x_smear.jpg. Smear images span from the day of puberty onset to the day of the last scan. In the growth_metrics folders, there are csv files containing daily body weight measurements, weekly body and tail length measurements.

Finally, the derivatives folder contains ANFNI_masks, AFNI_seed_connectivity, population_template, BEN_weight and RABIES folders. The RABIES folder contains important outputs from the rabies preprocessing and confound pipeline. Files include the confound corrected timeseries, temporal signal to noise (tSNR), framewise displacement (FD) and motion parameters. The AFNI_mask folder contains a strict mask for each confound-corrected timeseries file. The AFNI_seed_connectivity folder includes timeseries, r-correlation maps and z-correlation maps from all seeds used for technical validation. The population_template includes the population-average template at 0.2 and 0.3 mm resolution, along with all seed masks used for technical validation. The BEN_weight folder contains the custom weight used to generate the masks for each anatomical image prior to unbiased population template construction.

The dataset contains 180 anatomical scans and 179 functional scans. Due to technical issues during acquisition, sub-2023080406 does not have a ses-1 functional scan. Additionally, due to technical issues with physiological monitoring system, eight respiration data files were corrupted, so breathing data is unavailable for sub-2023080701_ses-3, sub-2023080702_ses-2, sub-2023080702_ses-3, sub-2023090101_ses-2, sub-2023090103_ses-3, sub-2023090106_ses-4, sub-2023090107_ses-4, sub-2024011506_ses-4

### Technical Validation

To assess the quality of the rs-fMRI scans, we computed a series of quality metrics previously developed to validate rat rs-fMRI datasets^36^ (https://github.com/grandjeanlab/MultiRat). First, we examined mean respiration during the functional scan for each session, as respiratory rate helps to estimate the depth of anaesthesia, with higher rates typically indicating superficial levels of anaesthesia and more robust functional connectivity^36^. Breathing rates during resting-state acquisition are comparable across development (see figure 2).

**Figure 2.**
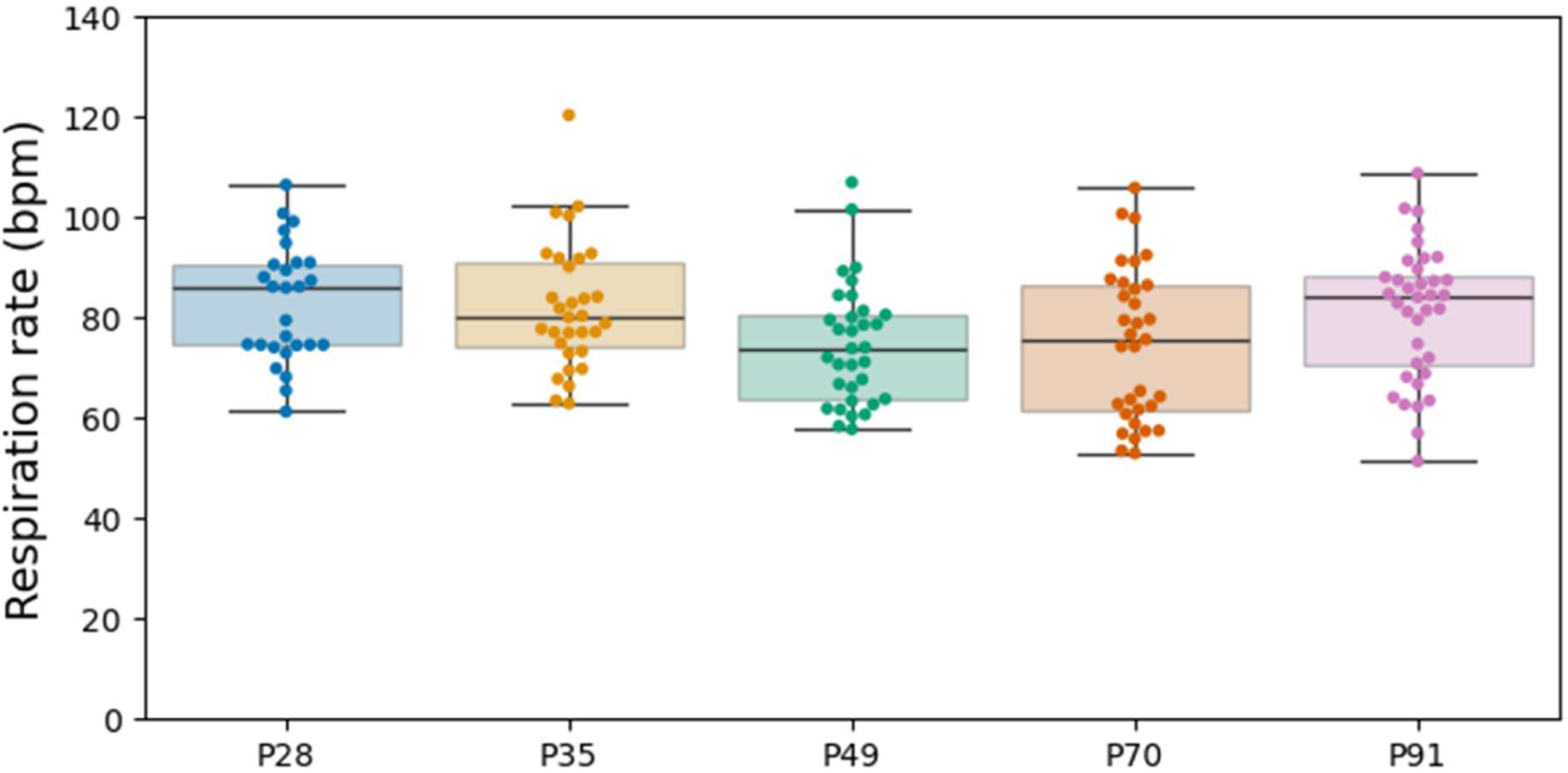
Mean respiration rate for each rat across five resting-state sessions. Each data point corresponds to the mean respiration rate for a rat during the resting-state scan for that timepoint. Boxplots show the median and interquartile range (IQR) for each timepoint. The lower whisker extends to the lowest data point above 1.5 x IQR. The upper whisker extends to the highest data point below 1.5 x IQR. Animals that failed preprocessing are excluded.

Next, we examined the degree of motion in each session, as high levels of head motion during a scan are a source of noise and artefacts, reducing data quality. We computed mean framewise displacement (FD; a measure of how much movement occurs from one frame to the next) for each session (see figure 3). During their first two development scans, rats displayed greater mean FD compared to later scans. This likely reflects the younger animals’ smaller size relative to the MRI cradle, which allowed for slightly more movement, relative to scans performed at older ages. Greater movement at younger ages is also a characteristic of human neuroimaging data^50,51^.

**Figure 3.**
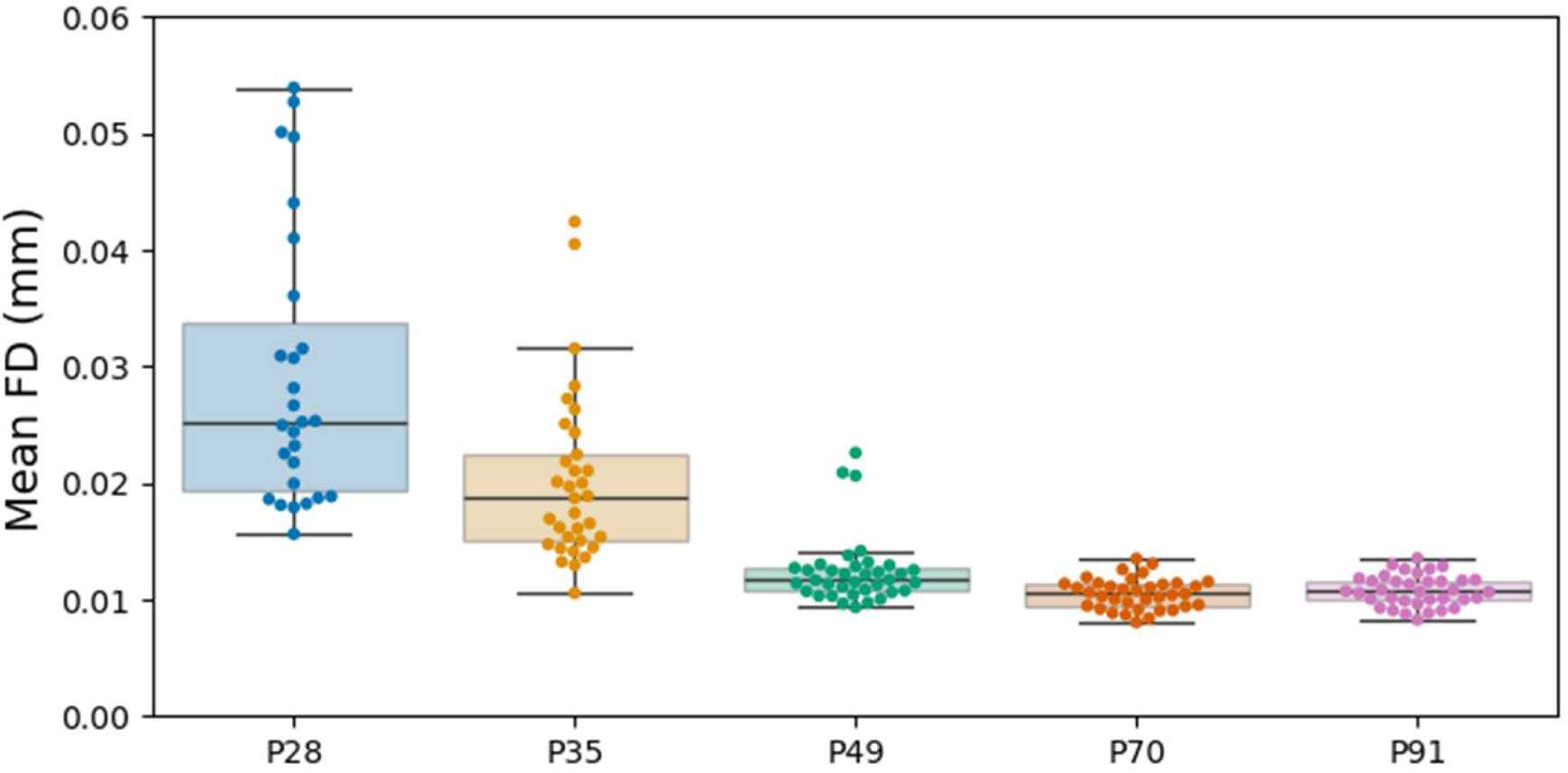
Mean framewise displacement for each scan across five resting-state sessions. Each data point corresponds to the mean framewise displacement for a rat during the resting-state scan for that timepoint. Boxplots show the median and interquartile range (IQR) for each timepoint. The lower whisker extends to the lowest data point above 1.5 x the IQR. The upper whisker extends to the highest data point below 1.5 x the IQR. Greater framewise displacement during Sessions 1 and 2 likely reflects the animals’ younger age and smaller size relative to the MRI cradle, which allowed for slightly more movement, compared to sessions 3-5, during which the animals were older and larger. Animals that failed preprocessing are excluded.

Next we examined the temporal signal-to-noise ratio (tSNR) using the tSNR maps generated by RABIES during preprocessing. tSNR provides a measure of how stable the fMRI signal is over time. The mean tSNR of the left and right barrel fields and left and right striatum were used to provide a cortical and subcortical measure of tSNR across sessions (see figure 4). While subcortical tSNR was similar across timepoints (age), the first two timepoints (P28, P35) collected had marginally lower cortical tSNR compared to later timepoints, which is likely related to the greater degree of movement during those juvenile age scans, but may also reflect developmental changes in the resting state signal.

**Figure 4.**
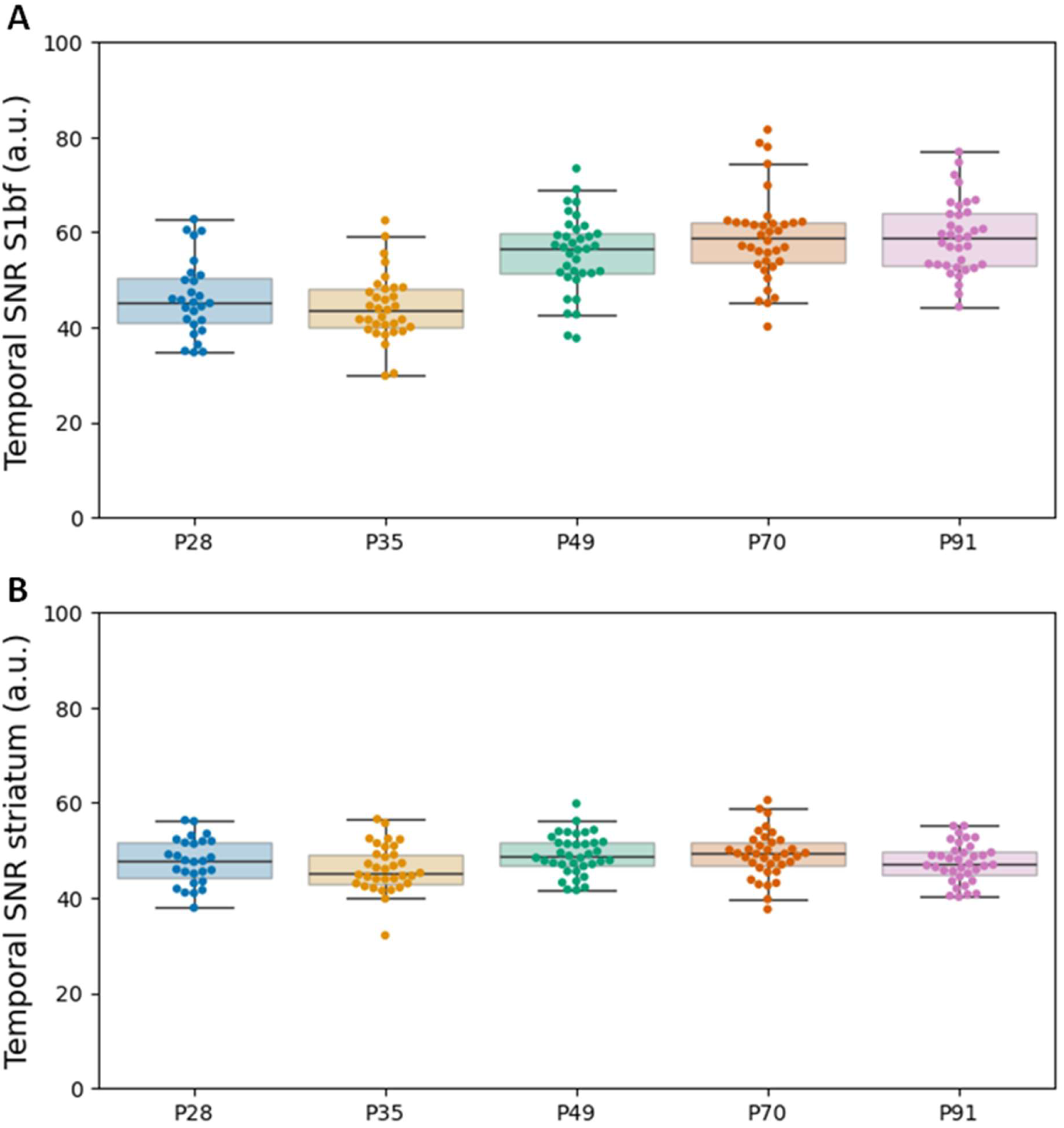
Cortical and subcortical temporal signal-to-noise for each scan across five resting-state sessions. Each data point corresponds to the mean temporal SNR for the left and right somatosensory barrel field (S1bf) or left and right striatum at each timepoint. The mean of the left and right S1bf was used as the cortical region (A) and the mean across the left and right striatum was used as the subcortical region (B). Boxplots show the median and interquartile range (IQR) for each timepoint. The lower whisker extends to the lowest data point above 1.5 x the IQR. The upper whisker extends to the highest data point below 1.5 x the IQR. Animals that failed preprocessing are excluded.

Finally, we carried out seed-based analysis to assess functional connectivity across sessions to enable comparison with existing rat datasets^36^. 0.9-mm diameter seeds located in the S1 barrel field (S1bf), anterior cingulate area (ACA), caudate-putamen and primary motor area previously used to validate the StandardRat protocol^36^ were transformed into our population-average space. To transform the seed ROIs, we first registered our template to the SIMGA rat atlas in vivo anatomical template v1.2.1^52^ and applied the inverse transform to the ROI seeds. The resulting ROIs were adjusted so that they retained their original size of 0.9mm. Individual timeseries and z-score correlations maps were generated using AFNI. Following StandardRat^36^, individual rat seed maps were thresholded at z > 1.96 (corresponding to p ≤ 0.05, one tailed), uncorrected. An uncorrected threshold is used to ensure that no FC is rejected (low false negative), but at the expense of a higher false-positive rate (see figure 6 **and 7**).

**Figure 5.**
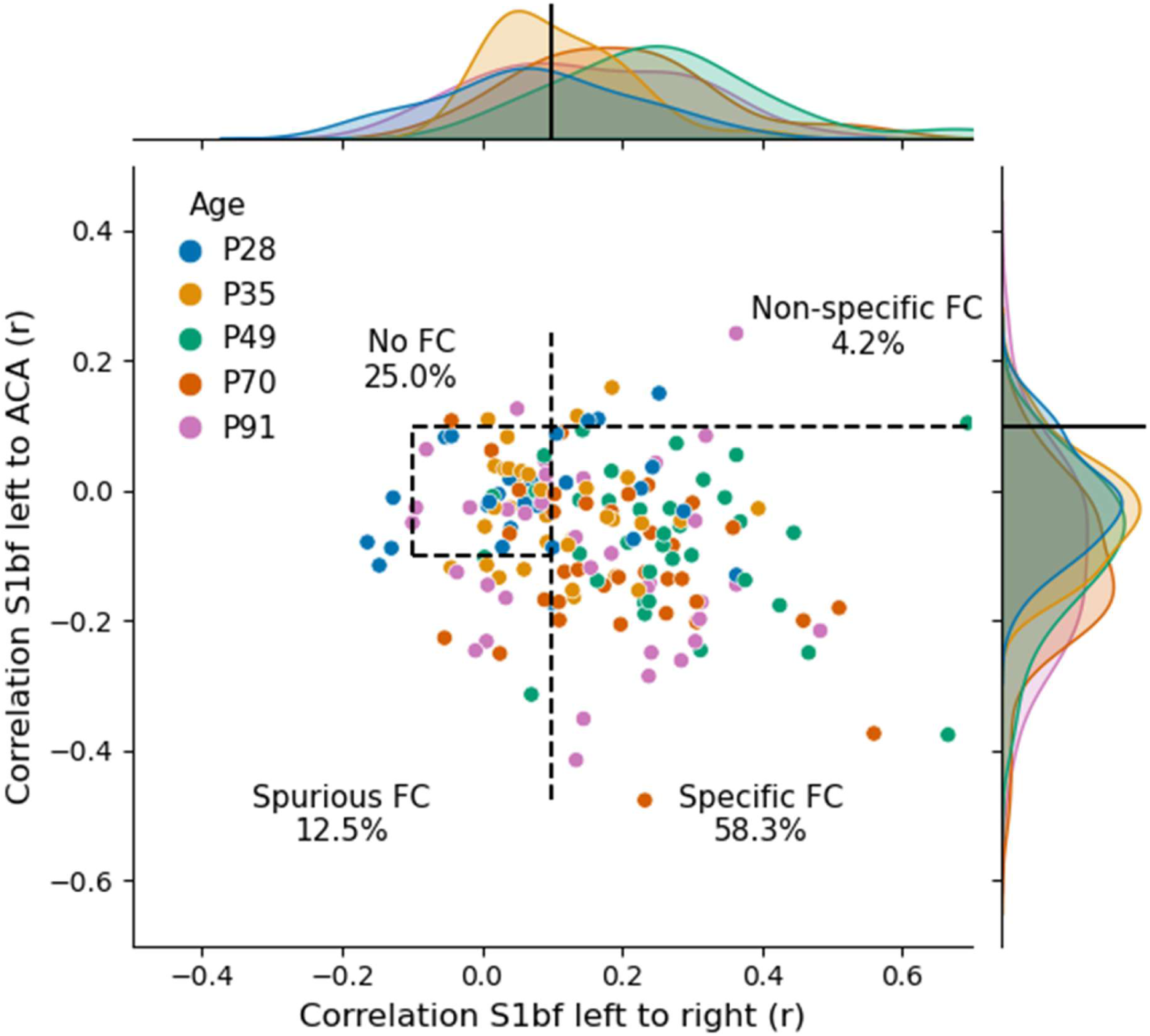
Categorising functional connectivity based on connectivity between specific and non-specific seeds. Functional connectivity in left somatosensory barrel field (S1bf) relative to **specific** (right S1bf) and to **non-specific** (anterior cingulate area - ACA) regions of interest. Each data point represents a scan; broken lines indicate the thresholds used to categorise each session into the four categories defined by Grandjean et al.^36^. Data points are coloured by session. **P28** (27 scans): specific (9), no FC (11), spurious (4), non-specific (3). **P35**: specific (13), no FC (13), spurious (5), non-specific (2). **P49**: specific (29), no FC (4), spurious (2), non-specific (1). **P70**: specific (28), no FC (4), spurious (4), non-specific (0). **P91**: specific FC (19), no FC (10), spurious (6), non-specific (1).

**Figure 6.**
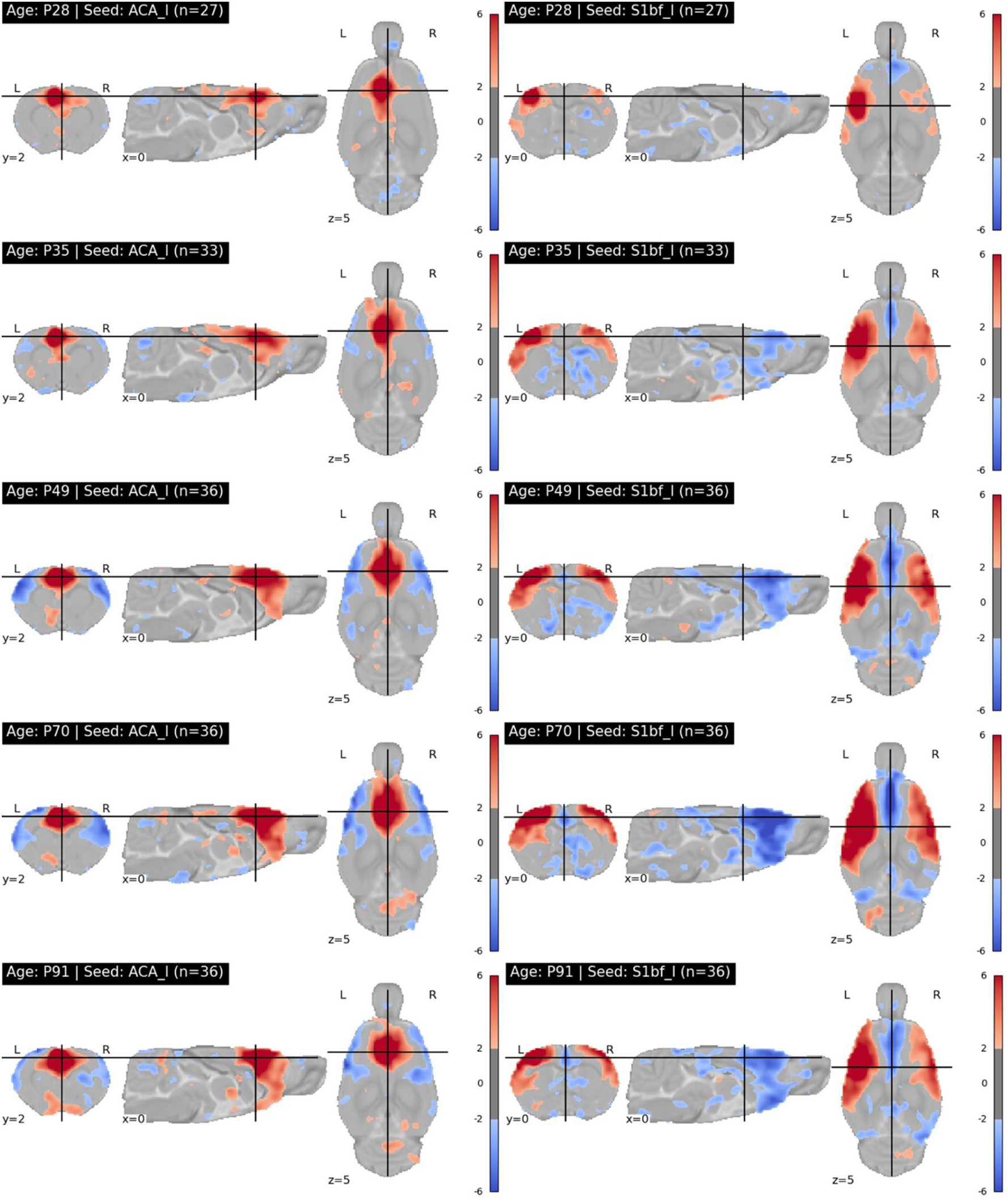
Correlation maps for seeds placed in the left left anterior cingulate area and left S1 barrel field. Spherical seeds (0.9mm diameter) were placed in the left anterior cingulate area (ACA_l) and left S1 barrel field (S1bf_l). Following the convention in Grandjean et al.^36^, for illustrative purposes, correlation maps were converted to z-scores and a maps and thresholded z > 1.96 corresponding to p ≤ 0.05, one tailed, uncorrected. This is to ensure that no FC is rejected (low false negative), but at the expense of a higher false-positive rate.

**Figure 7.**
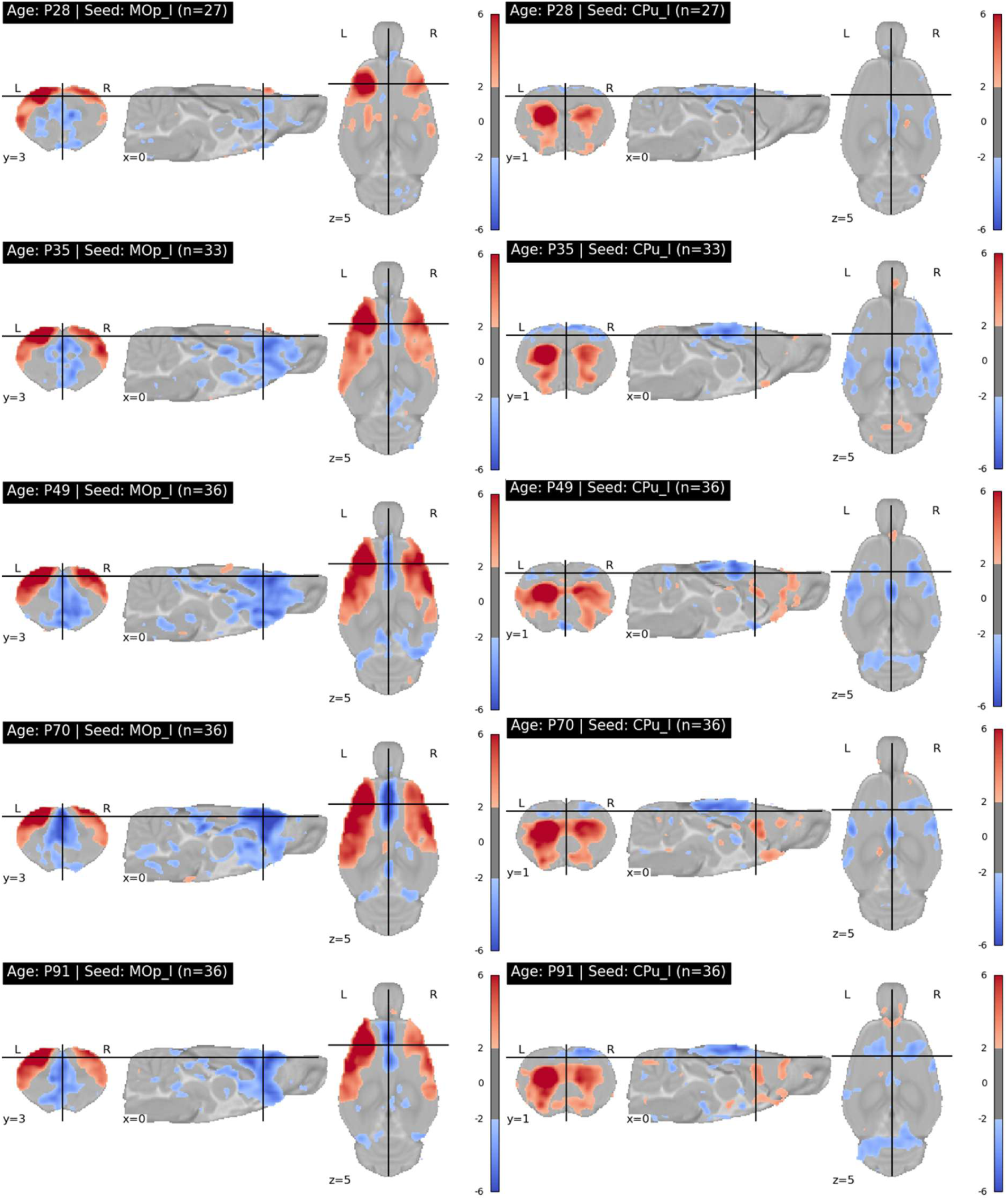
Correlation maps for seeds placed in the left primary motor cortex and left caudate-putamen. Spherical seeds (0.9mm diameter) were placed in the left primary motor cortex (MOp_l) and left caudate-putamen (CPu_l). For illustrative purposes, correlation maps were converted to z-score maps and thresholded z > 1.96 corresponding to p ≤ 0.05, one tailed, uncorrected. This is to ensure that no FC is rejected (low false negative), but at the expense of a higher false-positive rate.

Following the approach used in StandardRat^36^, we examined individual-level FC maps. Since sensory networks are particularly reliable for evaluating the effects of anaesthesia at depths commonly used in fMRI studies^53^, FC for sensory cortex was investigated. Using seeds placed in the left and right S1bf, we first looked for robust connectivity between inter-hemispheric sensory cortices, as most networks exhibit a bilateral homotopic organisation in both humans and animals^54^. Second, we investigated the correlation between the left S1bf seed and a seed positioned in the ACA, a key node in the rodent task-negative default mode network. Task-positive networks and task-negative networks are generally uncorrelated or anti-correlated, so a weak or negative correlation between these seeds is expected^55^.

Based on these expected patterns, and using mean FC computed between each of the three seeds (left S1bf, right S1bf, ACA) for each rat and each scan, we assigned scans into one of four categories: specific, non-specific, spurious and no connectivity. Specific FC was defined as a correlation between left and right S1bf of greater than 0.1, and a correlation between the left S1bf and ACA less than 0.1; non-specific was defined as a correlation between left and right S1bf of greater than 0.1, and a correlation between the left S1bf and ACA greater than 0.1; no connectivity was defined as a correlation between left and right S1bf ranging from -0.1 and 0.1, and a correlation between the left S1bf and ACA ranging from -0.1 and 0.1; and spurious was any remaining cases. Pearson r was used for all correlations. 58.3% of scans displayed specific FC, 12.5% displayed spurious FC, 25% displayed no FC and 4.2% had non-specific FC (see figure 5). A breakdown of assignments based on FC at each age is provided in **table 1**. These are in line with the results from the StandardRat consensus study: 61.8% of scans displayed specific FC, 3.9% displayed spurious FC, 28.5% displayed no FC and 5.8% had non-specific FC^36^. Inspection of figures 5, figure 6, and **table 1** provides some insight into developmental effects in these patterns: bilateral (interhemispheric) connectivity is weaker at the youngest ages (P28, P35), as is negative connectivity between sensory cortex and ACA.

**Table 1.** Summary of scan classification at each timepoint. Functional connectivity was computed between the left S1 barrel field (S1bf), right S1bf and left anterior cingulate area (ACA) for each scan. **Specific FC**: left S1bf to right S1bf *r* > 0.1 and left S1bf to ACA *r* < 0.1; **No FC**: left S1bf to right S1bf *r* = -0.1 to 0.1 and left S1bf to ACA *r* = -0.1 to 0.1; **Non-specific FC**: left S1bf to right S1bf *r* > 0.1 and left S1bf to ACA *r* > 0.1; **Spurious FC**: all other cases.

|  | P28 | P35 | P49 | P70 | P91 |
| --- | --- | --- | --- | --- | --- |
| Specific FC | 9 (33%) | 13 (39%) | 29 (81%) | 28 (78%) | 19 (53%) |
| No FC | 11 (41%) | 13 (39%) | 4 (11%) | 4 (11%) | 10 (28%) |
| Non-specific FC | 3 (11%) | 2 (6%) | 1 (3%) | 0 (0%) | 1 (3%) |
| Spurious FC | 4 (15%) | 5 (15%) | 2 (5%) | 4 (11%) | 6 (16%) |

### Usage Notes

All data are shared for reuse for non-commercial scientific research purposes. Users of the dataset are requested to cite this publication and OpenNeuro Dataset ds007859.

Users interested in further collaboration, particularly with respect to the linked postmortem brain bank, should contact the authors.

## Data Availability

All data are publicly available on OpenNeuro, Dataset ds007859, at the following link: https://openneuro.org/datasets/ds007859/

## Code Availability

Code to replicate the validation steps included here and run the entire template construction, preprocessing and confound correction pipeline is available on Codeberg at the following link: https://codeberg.org/mclooned/rat_normative_development_dataset

## Funding

This project was funded by Frontiers for the Future Grant 20/FFP-P/8799 to CK and AH from Taighde Éireann (Research Ireland).

## Author Contributions

D.M. was involved in study administration, animal handling, data acquisition, data analysis, technical validation, and writing of the manuscript.

A.B. was involved in study administration (not sure what this means so remove if I wasn’t), animal handling, data acquisition, and writing of the Introduction.

C.M. was involved in data acquisition and writing of the introduction.

A.H. obtained the funding and was involved in conceptualisation, and review of the manuscript.

C.K. obtained the funding and was involved in conceptualisation, study administration, review of the manuscript, and overall supervision of the study.

## Ethics declarations

### Competing Interests

The authors declare no competing interests.

### AI use

The authors declare that generative artificial intelligence has not been used for the writing of this manuscript, nor for the creation of figures, nor their corresponding legends.

## Notes

https://openneuro.org/datasets/ds007859/

### Competing Interest Statement

The authors have declared no competing interest.

## References

1. Rogol, A. D., Roemmich, J. N. & Clark, P. A. Growth at puberty. J. Adolesc. Health 31, 192–200 (2002).

2. Blakemore, S.-J. Adolescence and mental health. The Lancet 393, 2030–2031 (2019).

3. Spear, L. P. The adolescent brain and age-related behavioral manifestations. Neurosci. Biobehav. Rev. 24, 417–463 (2000).

4. Steinberg, L. Cognitive and affective development in adolescence. Trends Cogn. Sci. 9, 69–74 (2005).

5. Sisk, C. L. & Foster, D. L. The neural basis of puberty and adolescence. Nat. Neurosci. 7, 1040–1047 (2004).

6. Goddings, A.-L., Beltz, A., Peper, J. S., Crone, E. A. & Braams, B. R. Understanding the Role of Puberty in Structural and Functional Development of the Adolescent Brain. J. Res. Adolesc. 29, 32–53 (2019).

7. McEwen, B. S. & Milner, T. A. Understanding the broad influence of sex hormones and sex differences in the brain. J. Neurosci. Res. 95, 24–39 (2017).

8. Schulz, K. M. & Sisk, C. L. The organizing actions of adolescent gonadal steroid hormones on brain and behavioral development. Neurosci. Biobehav. Rev. 70, 148–158 (2016).

9. Saldanha, C. J. et al. Entanglement of Gender/Sex Dynamics in Basic and Developmental Systems Biology. in Sex and Gender: Toward Transforming Scientific Practice (eds DuBois, L. Z., Kaiser Trujillo, A. & McCarthy, M. M.) 23–47 (Springer Nature Switzerland, Cham, 2025). doi:10.1007/978-3-031-91371-6_2.

10. Andrews, J. L., Ahmed, S. P. & Blakemore, S.-J. Navigating the Social Environment in Adolescence: The Role of Social Brain Development. Biol. Psychiatry 89, 109–118 (2021).

11. Chaku, N. & Hoyt, L. T. Developmental Trajectories of Executive Functioning and Puberty in Boys and Girls. J. Youth Adolesc. 48, 1365–1378 (2019).

12. Galván, A. Adolescent Brain Development and Contextual Influences: A Decade in Review. J. Res. Adolesc. Off. J. Soc. Res. Adolesc. 31, 843–869 (2021).

13. Schulz, K. M. & Sisk, C. L. Pubertal hormones, the adolescent brain, and the maturation of social behaviors: Lessons from the Syrian hamster. Mol. Cell. Endocrinol. 254–255, 120–126 (2006).

14. Huttenlocher, P. R. Synaptic density in human frontal cortex — Developmental changes and effects of aging. Brain Res. 163, 195–205 (1979).

15. Wierenga, L. M. et al. Unraveling age, puberty and testosterone effects on subcortical brain development across adolescence. Psychoneuroendocrinology 91, 105–114 (2018).

16. Kessler, R. C. et al. Lifetime prevalence and age-of-onset distributions of mental disorders in the world health organization’s world mental health survey initiative. World Psychiatry 6, 168–176 (2007).

17. Andersen, S. L. Trajectories of brain development: Point of vulnerability or window of opportunity? Neurosci. Biobehav. Rev. 27, 3–18 (2003).

18. Andersen, S. L. & Teicher, M. H. Stress, sensitive periods and maturational events in adolescent depression. Trends Neurosci. 31, 183–191 (2008).

19. Tottenham, N. & Galván, A. Stress and the adolescent brain: Amygdala-prefrontal cortex circuitry and ventral striatum as developmental targets. Neurosci. Biobehav. Rev. 70, 217–227 (2016).

20. Angold, A. & Costello, E. J. Puberty and depression. Child Adolesc. Psychiatr. Clin. N. Am. 15, 919–937, ix (2006).

21. Hayward, C. & Sanborn, K. Puberty and the emergence of gender differences in psychopathology. J. Adolesc. Health Off. Publ. Soc. Adolesc. Med. 30, 49–58 (2002).

22. Patton, G. C. et al. Menarche and the onset of depression and anxiety in Victoria, Australia. J. Epidemiol. Community Health 50, 661–666 (1996).

23. Marquand, A. F. et al. Conceptualizing mental disorders as deviations from normative functioning. Mol. Psychiatry 24, 1415–1424 (2019).

24. Di Martino, A. et al. Unraveling the miswired connectome: a developmental perspective. Neuron 83, 1335–1353 (2014).

25. Bethlehem, R. a. I., et al. Brain charts for the human lifespan. Nature 604, 525–533 (2022).

26. Shafiei, G. et al. Reproducible Brain Charts: An open data resource for mapping brain development and its associations with mental health. Neuron 113, 3758–3779.e6 (2025).

27. Jernigan, T. L., Brown, S. A. & Dowling, G. J. The Adolescent Brain Cognitive Development Study. J. Res. Adolesc. Off. J. Soc. Res. Adolesc. 28, 154–156 (2018).

28. Fan, X.-R. et al. A longitudinal resource for population neuroscience of school-age children and adolescents in China. Sci. Data 10, 545 (2023).

29. Liu, S. et al. Chinese Color Nest Project : An accelerated longitudinal brain-mind cohort. Dev. Cogn. Neurosci. 52, 101020 (2021).

30. Zhou, Z.-X., Chen, L.-Z., Milham, M. P. & Zuo, X.-N. Six cornerstones for translational brain charts. Sci. Bull. 68, 795–799 (2023).

31. Louis, T. A., Robins, J., Dockery, D. W., Spiro, A. & Ware, J. H. Explaining discrepancies between longitudinal and cross-sectional models. J. Chronic Dis. 39, 831–839 (1986).

32. Telzer, E. H. et al. Methodological considerations for developmental longitudinal fMRI research. Dev. Cogn. Neurosci. 33, 149–160 (2018).

33. Bettencourt, L. M. A., Stephens, G. J., Ham, M. I. & Gross, G. W. Functional structure of cortical neuronal networks grown in vitro. Phys. Rev. E Stat. Nonlin. Soft Matter Phys. 75, 021915 (2007).

34. Lu, H. et al. Rat brains also have a default mode network. Proc. Natl. Acad. Sci. U. S. A. 109, 3979–84 (2012).

35. Schwarz, A. J., Gozzi, A. & Bifone, A. Community structure and modularity in networks of correlated brain activity. Magn. Reson. Imaging 26, 914–920 (2008).

36. Grandjean, J. et al. A consensus protocol for functional connectivity analysis in the rat brain. Nat. Neurosci. 26, 673–681 (2023).

37. Shansky, R. M. & Woolley, C. S. Considering sex as a biological variable will be valuable for neuroscience research. J. Neurosci. 36, 11817–11822 (2016).

38. Will, T. R. et al. Problems and progress regarding sex bias and omission in neuroscience research. eNeuro 4, ENEURO.0278-17.2017 (2017).

39. Becker, J. B., Prendergast, B. J. & Liang, J. W. Female rats are not more variable than male rats: a meta-analysis of neuroscience studies. Biol. Sex Differ. 7, 34 (2016).

40. Dalla, C. et al. Practical solutions for including sex as a biological variable (SABV) in preclinical neuropsychopharmacological research. J. Neurosci. Methods 401, 110003 (2024).

41. Shansky, R. M. Behavioral neuroscience’s inevitable SABV growing pains. Trends Neurosci. 47, 669–676 (2024).

42. McCarthy, M. M., Woolley, C. S. & Arnold, A. P. Incorporating sex as a biological variable in neuroscience: What do we gain? Nat. Rev. Neurosci. 18, 707–708 (2017).

43. Lauwereyns, J. et al. Toward a common interpretation of the 3Rs principles in animal research. Lab Anim. 53, 347–350 (2024).

44. Gaytan, F. et al. Development and validation of a method for precise dating of female puberty in laboratory rodents: The puberty ovarian maturation score (Pub-Score). Sci. Rep. 7, 46381 (2017).

45. Gaytan, F., Bellido, C., Aguilar, R. & Aguilar, E. Balano-preputial separation as an external sign of puberty in the rat: correlation with histologic testicular data. Andrologia 20, 450–453 (1988).

46. Vrooman, R. M. et al. fMRI data acquisition and analysis for task-free, anesthetized rats. Nat. Protoc. 20, 1393–1412 (2025).

47. Yu, Z. et al. A generalizable brain extraction net (BEN) for multimodal MRI data from rodents, nonhuman primates, and humans. eLife 11, e81217 (2022).

48. Germann, J., Gouveia, F. V., Chakravarty, M. M. & Devenyi, G. A. Longitudinal deformation-based morphometry pipeline to study neuroanatomical differences in structural MRI based on SyN unbiased templates. Aperture Neuro 5, (2025).

49. Desrosiers-Grégoire, G., Devenyi, G. A., Grandjean, J. & Chakravarty, M. M. A standardized image processing and data quality platform for rodent fMRI. Nat. Commun. 15, 6708 (2024).

50. Van Dijk, K. R. A., Sabuncu, M. R. & Buckner, R. L. The influence of head motion on intrinsic functional connectivity MRI. NeuroImage 59, 431–438 (2012).

51. Yan, C.-G. et al. A comprehensive assessment of regional variation in the impact of head micromovements on functional connectomics. NeuroImage 76, 183–201 (2013).

52. Barrière, D. A. et al. The SIGMA rat brain templates and atlases for multimodal MRI data analysis and visualization. Nat. Commun. 10, 5699 (2019).

53. Paasonen, J., Stenroos, P., Salo, R. A., Kiviniemi, V. & Gröhn, O. Functional connectivity under six anesthesia protocols and the awake condition in rat brain. NeuroImage 172, 9–20 (2018).

54. Biswal, B., Yetkin, F. Z., Haughton, V. M. & Hyde, J. S. Functional connectivity in the motor cortex of resting human brain using echo-planar MRI. Magn. Reson. Med. 34, 537–541 (1995).

55. Fox, M. D. et al. The human brain is intrinsically organized into dynamic, anticorrelated functional networks. Proc. Natl. Acad. Sci. U. S. A. 102, 9673–9678 (2005).

